## Supplementary Materials for "More rule than exception: Parallel evidence of ancient migrations in grammars and genomes of Finno-Ugric speakers"

**Supplementary Material**.

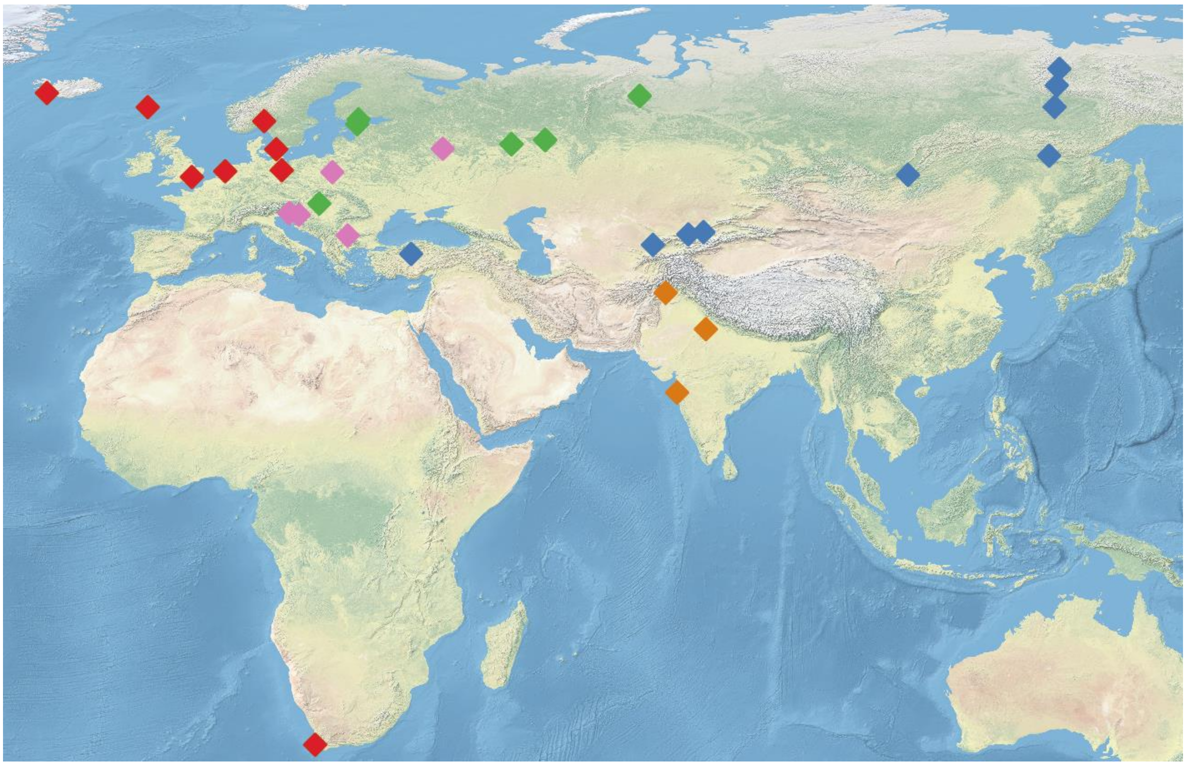

**Supplementary Figure S1.** Approximate geographical location of the 34 languages considered. Thirty-one geographical points; three FU languages, Mari, Udmurt and Khanty, are represented each by two diastratic variants spoken in the same location. Language groups coded as follows: Finno-Ugric (green); Altaic (blue); Germanic (red); Slavic (pink); Indo-Iranian (orange).

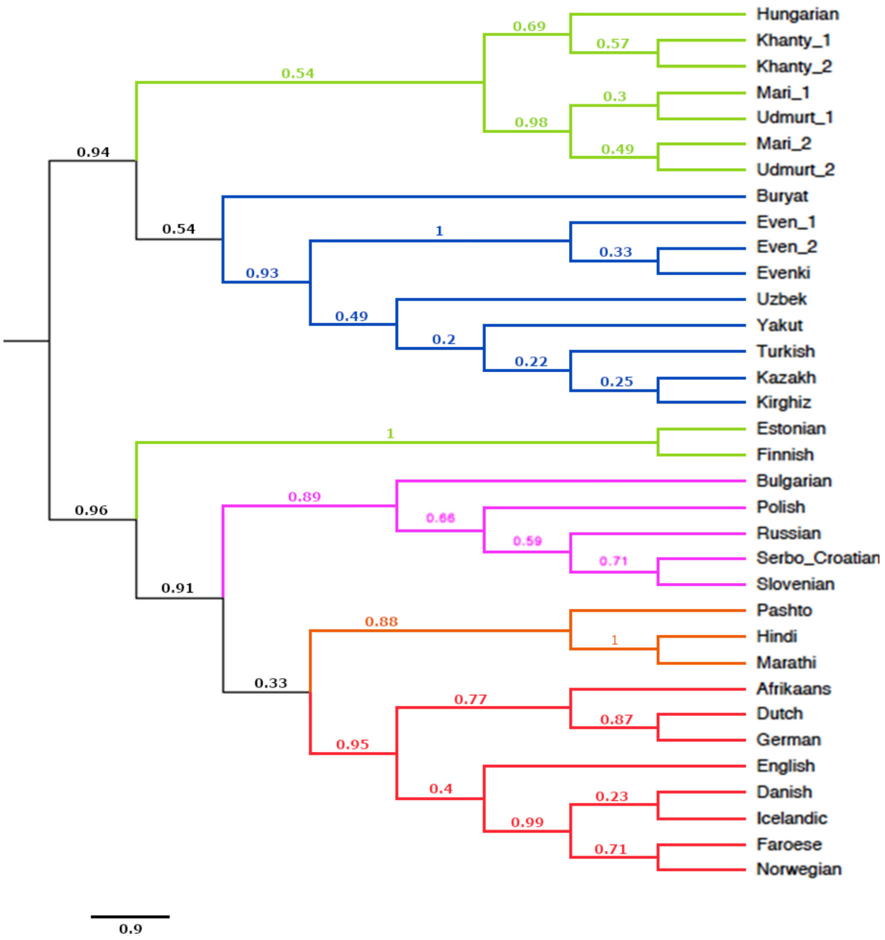


**Supplementary Figure S2.** Bayesian phylogeny (BEAST) from the syntactic dataset. Numbers on the nodes represent the posterior probability. From the top: Yellow= Altaic, light brown=Finno-Ugric, purple=Slavic, blue=Indo-Iranian, red=Germanic.

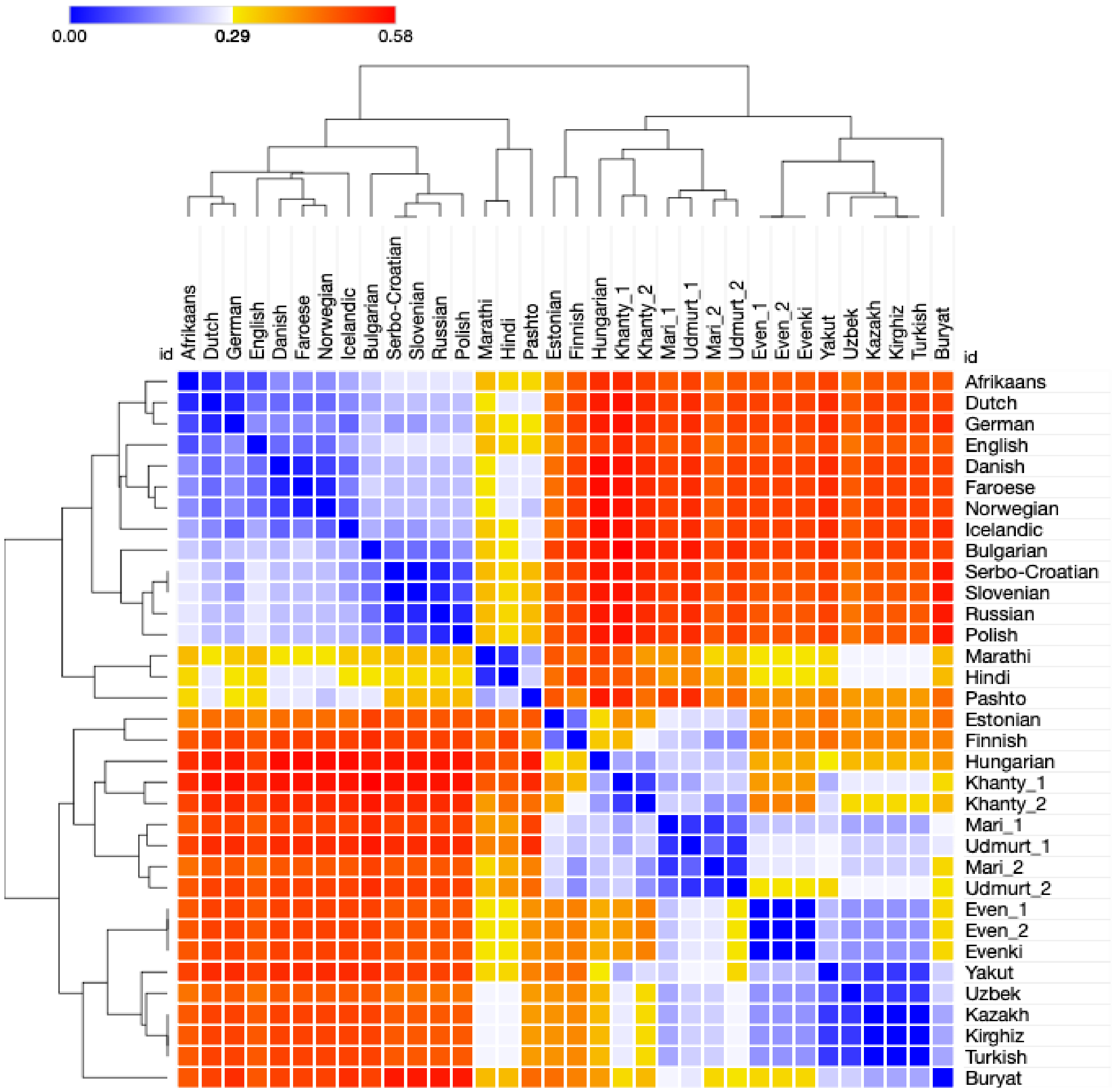


**Supplementary Figure S3.** Heatmap from the syntactic distances. Dark red represents maximum distance, dark blue minimum distance.

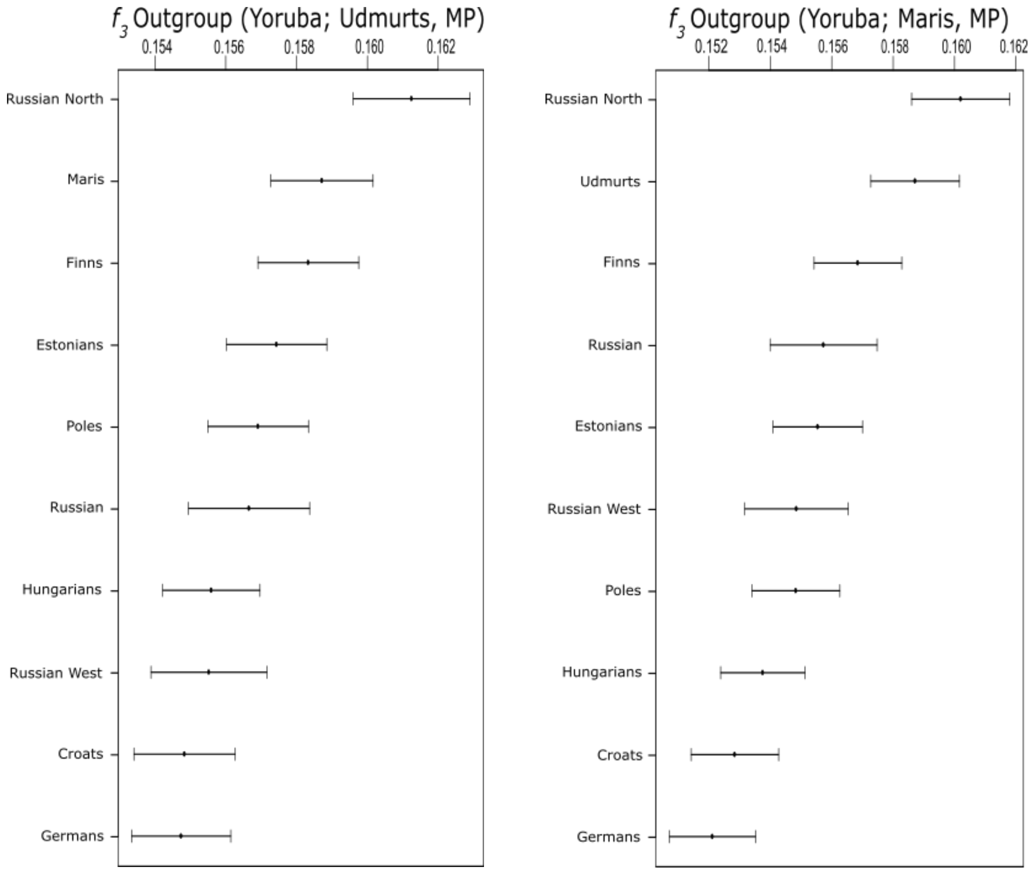


**Supplementary Figure S4.** Outgroup f3-statistics analysis. Shared genetic drift between modern Pontic steppes populations and modern European populations (MP).

**Supplementary Table S1.** Whole-genome samples collected for the populations under study.

| Sample Size | Sample ID | Populations | Country | Region | Coverage | Language Family | Reference |
|---|---|---|---|---|---|---|---|
| 3 | Est1, Est2, Est3 | **Estonian** | Estonia | Europe | >40 | **Uralic** | Pagani *et al.* 2016 |
| 3 | Fin1, Fin2, Fin3 | **Finnish** | Finland | Europe | >40 | **Uralic** | Pagani *et al.* 2016 |
| 3 | Rus1, RusPi1, RusPs2 | **Russian (North and West)** | Russia | Europe | >40 | **Indo-European** | Pagani *et al.* 2016 |
| 3 | Pole1, Pole2, Pole3 | **Polish** | Poland | Europe | >40 | **Indo-European** | Pagani *et al.* 2016 |
| 3 | Hun1, Hun2, Hun4 | **Hungarian** | Hungary | Europe | >40 | **Uralic** | Pagani *et al.* 2016 |
| 3 | Ger1, Ger2, Ger3 | **German** | Germany | Europe | >40 | **Indo-European** | Pagani *et al.* 2016 |
| 3 | croat11, croat13, croat12 | **Croatian** | Bosnia-Herzegovina | Europe | >40 | **Indo-European** | Pagani *et al.* 2016 |
| 3 | Iran1, Iran2, Iran3 | **Iranian (Farsi)** | Iran | West Asia | >40 | **Indo-European** | Pagani *et al.* 2016 |
| 3 | Mari1, Mari2, Mari3 | **Mari** | Russia | Europe | >40 | **Uralic** | Pagani *et al.* 2016 |
| 3 | Udmrd1, Udmrd2, Udmrd3 | **Udmurt** | Russia | Europe | >40 | **Uralic** | Pagani *et al.* 2016 |
| 3 | Khant1, Khant2, Khant3 | **Khanty** | Russia | Siberia | >40 | **Uralic** | Pagani *et al.* 2016 |
| 3 | Evnk2, Evk14, Evk16 | **Evenki** | Russia | Siberia | >40 | **Altaic** | Pagani *et al.* 2016 |
| 3 | Bur2, Bur6, Bur11 | **Buryat** | Russia | Siberia | >40 | **Altaic** | Pagani *et al.* 2016 |
| 3 | YakS4, YakK1, YakK3 | **Yakut** | Russia | Siberia | >40 | **Altaic** | Pagani *et al.* 2016 |
| 3 | EvenM1, EvenM2, EvenM3 | **Even** | Russia | Siberia | >40 | **Altaic** | Pagani *et al.* 2016 |

Note: the three Russian individuals come from three different subsets.

**Supplementary Table S2.** Ancient DNA samples used in this study.

| Sample ID | Archaeological Culture | Date | Country | Region | Contributor |
|---|---|---|---|---|---|
| I1100 | Anatolia_Neolithic | 6500-6200 calBCE | Turkey | Barcın | Mathieson *et al.* 2015 |
| I1102 | Anatolia_Neolithic | 6500-6200 calBCE | Turkey | Barcın | Mathieson *et al.* 2015 |
| I1099 | Anatolia_Neolithic | 6500-6200 calBCE | Turkey | Barcın | Mathieson *et al.* 2015 |
| I1103 | Anatolia_Neolithic | 6400-5600 calBCE | Turkey | Barcın | Mathieson *et al.* 2015 |
| I1101 | Anatolia_Neolithic | 6400-5600 calBCE | Turkey | Barcın | Mathieson *et al.* 2015 |
| I1097 | Anatolia_Neolithic | 6400-5600 calBCE | Turkey | Barcın | Mathieson *et al.* 2015 |
| I0744 | Anatolia_Neolithic | 6400-5600 calBCE | Turkey | Barcın | Mathieson *et al.* 2015 |
| I1579 | Anatolia_Neolithic | 6500-6200 calBCE | Turkey | Barcın | Mathieson *et al.* 2015 |
| I1581 | Anatolia_Neolithic | 6500-6200 calBCE | Turkey | Barcın | Mathieson *et al.* 2015 |
| I1096 | Anatolia_Neolithic | 6500-6200 calBCE | Turkey | Barcın | Mathieson *et al.* 2015 |
| I1580 | Anatolia_Neolithic | 6500-6200 calBCE | Turkey | Barcın | Mathieson *et al.* 2015 |
| I1098 | Anatolia_Neolithic | 6500-6200 calBCE | Turkey | Barcın | Mathieson *et al.* 2015 |
| I1585 | Anatolia_Neolithic | 6500-6200 calBCE | Turkey | Barcın | Mathieson *et al.* 2015 |
| I0708 | Anatolia_Neolithic | 6500-6200 calBCE | Turkey | Barcın | Mathieson *et al.* 2015 |
| I0745 | Anatolia_Neolithic | 6500-6200 calBCE | Turkey | Barcın | Mathieson *et al.* 2015 |
| I0746 | Anatolia_Neolithic | 6500-6200 calBCE | Turkey | Barcın | Mathieson *et al.* 2015 |
| I1583 | Anatolia_Neolithic | 6500-6200 calBCE | Turkey | Barcın | Mathieson *et al.* 2015 |
| I0707 | Anatolia_Neolithic | 6500-6200 calBCE | Turkey | Barcın | Mathieson *et al.* 2015 |
| I0709 | Anatolia_Neolithic | 6500-6200 calBCE | Turkey | Barcın | Mathieson *et al.* 2015 |
| I0725 | Anatolia_Neolithic | 6500-6200 calBCE | Turkey | Barcın | Mathieson *et al.* 2015 |
| I0727 | Anatolia_Neolithic | 6500-6200 calBCE | Turkey | Barcın | Mathieson *et al.* 2015 |
| I0724 | Anatolia_Neolithic | 6500-6200 calBCE | Turkey | Barcın | Mathieson *et al.* 2015 |
| I0736 | Anatolia_Neolithic | 6500-6200 calBCE | Turkey | Barcın | Mathieson *et al.* 2015 |

| | | | | | |
|---|---|---|---|---|---|
| I0723 | Anatolia_Neolithic | 6500-6200 calBCE | Turkey | Barcın | Mathieson *et al.* 2015 |
| I0726 | Anatolia_Neolithic | 6500-6200 calBCE | Turkey | Barcın | Mathieson *et al.* 2015 |
| I0231 | Yamnaya | 2910-2875 calBCE | Russia | Ekaterinovka, Southern Steppe, Samara | Haak *et al.* 2015 |
| I0357 | Yamnaya | 3090-2910 calBCE | Russia | Lopatino I, Sok River, Samara | Haak *et al.* 2015 |
| I0370 | Yamnaya | 3500-2700 calBCE | Russia | Ishkinovka I, Eastern Orenburg, Pre-Ural steppe, Samara | Haak *et al.* 2015 |
| I0429 | Yamnaya | 3339-2917 calBCE | Russia | Lopatino I, Sok River, Samara | Haak *et al.* 2015 |
| I0438 | Yamnaya | 3021-2635 calBCE | Russia | Luzkhi I, Samara River, Samara | Haak *et al.* 2015 |
| I0439 | Yamnaya | 3305-2925 calBCE | Russia | Lopatino I, Sok River, Samara | Haak *et al.* 2015 |
| I0441 | Yamnaya | 3010-2622 calBCE | Russia | Kurmanaevka III, Buzuluk, Samara | Haak *et al.* 2015 |
| I0443 | Yamnaya | 3335-2912 calBCE | Russia | Grachevka II, Sok_River, Samara | Haak *et al.* 2015 |
| I0444 | Yamnaya | 3335-2881 calBCE | Russia | Kutuluk I, Kutuluk River, Samara | Haak *et al.* 2015 |
| RISE386 | Sintashta | 2298-2045 calBCE | Russia | Bulanovo | Allentoft *et al.* 2015 |
| RISE391 | Sintashta | 2120-1887 calBCE | Kazakhstan | Tanabergen II | Allentoft *et al.* 2015 |
| RISE392 | Sintashta | 2126-1896 calBCE | Russia | Stepnoe VII | Allentoft *et al.* 2015 |
| RISE394 | Sintashta | 1949-1754 calBCE | Russia | Bulanovo | Allentoft *et al.* 2015 |
| RISE395 | Sintashta | 1960-1756 calBCE | Russia | Bol'shekaraganskii | Allentoft *et al.* 2015 |

**Supplementary Table S3.** Human Origins data on present-day humans used in this study.

| Sample ID | Population | Country | Region | Contributor |
|---|---|---|---|---|
| Nov_005 | Nganasan | Russia | Central Asia Siberia | Lazaridis *et al.* 2014 |
| ADR00514 | Nganasan | Russia | Central Asia Siberia | Lazaridis *et al.* 2014 |
| ADR00513 | Nganasan | Russia | Central Asia Siberia | Lazaridis *et al.* 2014 |
| ADR00509 | Nganasan | Russia | Central Asia Siberia | Lazaridis *et al.* 2014 |
| ADR00512 | Nganasan | Russia | Central Asia Siberia | Lazaridis *et al.* 2014 |
| ADR00504 | Nganasan | Russia | Central Asia Siberia | Lazaridis *et al.* 2014 |
| ADR00507 | Nganasan | Russia | Central Asia Siberia | Lazaridis *et al.* 2014 |
| ADR00511 | Nganasan | Russia | Central Asia Siberia | Lazaridis *et al.* 2014 |
| ADR00510 | Nganasan | Russia | Central Asia Siberia | Lazaridis *et al.* 2014 |
| ADR00508 | Nganasan | Russia | Central Asia Siberia | Lazaridis *et al.* 2014 |
| ADR00515 | Nganasan | Russia | Central Asia Siberia | Lazaridis *et al.* 2014 |
| HGDP00774 | Han | China | East Asia | Patterson *et al.* 2012 |
| HGDP00775 | Han | China | East Asia | Patterson *et al.* 2012 |
| HGDP00776 | Han | China | East Asia | Patterson *et al.* 2012 |
| HGDP00777 | Han | China | East Asia | Patterson *et al.* 2012 |
| HGDP00779 | Han | China | East Asia | Patterson *et al.* 2012 |
| HGDP00780 | Han | China | East Asia | Patterson *et al.* 2012 |
| HGDP00781 | Han | China | East Asia | Patterson *et al.* 2012 |
| HGDP00782 | Han | China | East Asia | Patterson *et al.* 2012 |
| HGDP00783 | Han | China | East Asia | Patterson *et al.* 2012 |
| HGDP00784 | Han | China | East Asia | Patterson *et al.* 2012 |
| HGDP00785 | Han | China | East Asia | Patterson *et al.* 2012 |
| HGDP00786 | Han | China | East Asia | Patterson *et al.* 2012 |
| HGDP00811 | Han | China | East Asia | Patterson *et al.* 2012 |
| HGDP00812 | Han | China | East Asia | Patterson *et al.* 2012 |
| HGDP00813 | Han | China | East Asia | Patterson *et al.* 2012 |
| HGDP00814 | Han | China | East Asia | Patterson *et al.* 2012 |
| HGDP00815 | Han | China | East Asia | Patterson *et al.* 2012 |
| HGDP00817 | Han | China | East Asia | Patterson *et al.* 2012 |
| HGDP00818 | Han | China | East Asia | Patterson *et al.* 2012 |
| HGDP00819 | Han | China | East Asia | Patterson *et al.* 2012 |
| HGDP00820 | Han | China | East Asia | Patterson *et al.* 2012 |
| HGDP00821 | Han | China | East Asia | Patterson *et al.* 2012 |
| HGDP00822 | Han | China | East Asia | Patterson *et al.* 2012 |
| HGDP00971 | Han | China | East Asia | Patterson *et al.* 2012 |
| HGDP00972 | Han | China | East Asia | Patterson *et al.* 2012 |
| HGDP00973 | Han | China | East Asia | Patterson *et al.* 2012 |
| HGDP00974 | Han | China | East Asia | Patterson *et al.* 2012 |

| | | | | |
|---|---|---|---|---|
| HGDP00975 | Han | China | East Asia | Patterson *et al.* 2012 |
| HGDP00976 | Han | China | East Asia | Patterson *et al.* 2012 |
| HGDP00977 | Han | China | East Asia | Patterson *et al.* 2012 |
| HGDP01021 | Han | China | East Asia | Patterson *et al.* 2012 |
| HGDP01023 | Han | China | East Asia | Patterson *et al.* 2012 |
| HGDP01024 | Han | China | East Asia | Patterson *et al.* 2012 |
| HGDP00449 | Mbuti | Congo | Africa | Patterson *et al.* 2012 |
| HGDP00462 | Mbuti | Congo | Africa | Patterson *et al.* 2012 |
| HGDP00463 | Mbuti | Congo | Africa | Patterson *et al.* 2012 |
| HGDP00467 | Mbuti | Congo | Africa | Patterson *et al.* 2012 |
| HGDP00474 | Mbuti | Congo | Africa | Patterson *et al.* 2012 |
| HGDP00476 | Mbuti | Congo | Africa | Patterson *et al.* 2012 |
| HGDP00478 | Mbuti | Congo | Africa | Patterson *et al.* 2012 |
| HGDP00982 | Mbuti | Congo | Africa | Patterson *et al.* 2012 |
| HGDP00984 | Mbuti | Congo | Africa | Patterson *et al.* 2012 |
| HGDP01081 | Mbuti | Congo | Africa | Patterson *et al.* 2012 |
| HGDP00995 | Karitiana | Brazil | America | Patterson *et al.* 2012 |
| HGDP00999 | Karitiana | Brazil | America | Patterson *et al.* 2012 |
| HGDP01001 | Karitiana | Brazil | America | Patterson *et al.* 2012 |
| HGDP01003 | Karitiana | Brazil | America | Patterson *et al.* 2012 |
| HGDP01006 | Karitiana | Brazil | America | Patterson *et al.* 2012 |
| HGDP01010 | Karitiana | Brazil | America | Patterson *et al.* 2012 |
| HGDP01012 | Karitiana | Brazil | America | Patterson *et al.* 2012 |
| HGDP01013 | Karitiana | Brazil | America | Patterson *et al.* 2012 |
| HGDP01014 | Karitiana | Brazil | America | Patterson *et al.* 2012 |
| HGDP01015 | Karitiana | Brazil | America | Patterson *et al.* 2012 |
| HGDP01018 | Karitiana | Brazil | America | Patterson *et al.* 2012 |
| HGDP01019 | Karitiana | Brazil | America | Patterson *et al.* 2012 |
| Ul5 | Ulchi | Russia | Central Asia Siberia | Rem Sukernik / Stanislav Dryomov |
| Ul31 | Ulchi | Russia | Central Asia Siberia | Rem Sukernik / Stanislav Dryomov |
| Ul65 | Ulchi | Russia | Central Asia Siberia | Rem Sukernik / Stanislav Dryomov |
| Ul6 | Ulchi | Russia | Central Asia Siberia | Rem Sukernik / Stanislav Dryomov |
| Ul33 | Ulchi | Russia | Central Asia Siberia | Rem Sukernik / Stanislav Dryomov |
| Ul71 | Ulchi | Russia | Central Asia Siberia | Rem Sukernik / Stanislav Dryomov |
| Ul10 | Ulchi | Russia | Central Asia Siberia | Rem Sukernik / Stanislav Dryomov |
| Ul43 | Ulchi | Russia | Central Asia Siberia | Rem Sukernik / Stanislav Dryomov |
| Ul72 | Ulchi | Russia | Central Asia Siberia | Rem Sukernik / Stanislav Dryomov |
| Ul44 | Ulchi | Russia | Central Asia Siberia | Rem Sukernik / Stanislav Dryomov |
| Ul74 | Ulchi | Russia | Central Asia Siberia | Rem Sukernik / Stanislav Dryomov |
| Ul19 | Ulchi | Russia | Central Asia Siberia | Rem Sukernik / Stanislav Dryomov |
| Ul24 | Ulchi | Russia | Central Asia Siberia | Rem Sukernik / Stanislav Dryomov |

| | | | | |
|---|---|---|---|---|
| Ul59 | Ulchi | Russia | Central Asia Siberia | Rem Sukernik / Stanislav Dryomov |
| Ul56 | Ulchi | Russia | Central Asia Siberia | Rem Sukernik / Stanislav Dryomov |
| Ul55 | Ulchi | Russia | Central Asia Siberia | Rem Sukernik / Stanislav Dryomov |
| Ul16 | Ulchi | Russia | Central Asia Siberia | Rem Sukernik / Stanislav Dryomov |
| Ul69 | Ulchi | Russia | Central Asia Siberia | Rem Sukernik / Stanislav Dryomov |
| Ul1 | Ulchi | Russia | Central Asia Siberia | Rem Sukernik / Stanislav Dryomov |
| Ul36 | Ulchi | Russia | Central Asia Siberia | Rem Sukernik / Stanislav Dryomov |
| Ul25 | Ulchi | Russia | Central Asia Siberia | Rem Sukernik / Stanislav Dryomov |
| Ul52 | Ulchi | Russia | Central Asia Siberia | Rem Sukernik / Stanislav Dryomov |
| Ul70 | Ulchi | Russia | Central Asia Siberia | Rem Sukernik / Stanislav Dryomov |
| Ul51 | Ulchi | Russia | Central Asia Siberia | Rem Sukernik / Stanislav Dryomov |
| Ul39 | Ulchi | Russia | Central Asia Siberia | Rem Sukernik / Stanislav Dryomov |
| mixe0029 | Mixe | Mexico | America | William Klitz / Cheryl Winkle |
| mixe0030 | Mixe | Mexico | America | William Klitz / Cheryl Winkle |
| mixe0015 | Mixe | Mexico | America | William Klitz / Cheryl Winkle |
| mixe0035 | Mixe | Mexico | America | William Klitz / Cheryl Winkle |
| mixe0018 | Mixe | Mexico | America | William Klitz / Cheryl Winkle |
| mixe0026 | Mixe | Mexico | America | William Klitz / Cheryl Winkle |
| mixe0027 | Mixe | Mexico | America | William Klitz / Cheryl Winkle |
| mixe0028 | Mixe | Mexico | America | William Klitz / Cheryl Winkle |
| mixe0007 | Mixe | Mexico | America | William Klitz / Cheryl Winkle |
| mixe0009 | Mixe | Mexico | America | William Klitz / Cheryl Winkle |

**Supplementary Table S4.** Statistics of the *qpAdm* models.

| Test | Outgroup set | Nganasan | Yamnaya | Anatolia | chi-square |
|---|---|---|---|---|---|
| Khanty | Han, Mbuti, Karitiana, Ulchi and Mixe | 0.521 | 0.479 | 0 | 10.056 |
| Maris | Han, Mbuti, Karitiana, Ulchi and Mixe | 0.281 | 0.465 | 0.254 | 10.493 |
| Udmurts | Han, Mbuti, Karitiana, Ulchi and Mixe | 0.261 | 0.611 | 0.128 | 9.032 |
| Iranians | Han, Mbuti, Karitiana, Ulchi and Mixe | 0.016 | 0.141 | 0.843 | 7.053 |
| Finns | Han, Mbuti, Karitiana, Ulchi and Mixe | 0.101 | 0.589 | 0.31 | 6.623 |
| Estonians | Han, Mbuti, Karitiana, Ulchi and Mixe | 0.04 | 0.568 | 0.391 | 8.095 |
| Hungarians | Han, Mbuti, Karitiana, Ulchi and Mixe | 0.032 | 0.412 | 0.556 | 8.398 |
| Russian North | Han, Mbuti, Karitiana, Ulchi and Mixe | 0.144 | 0.571 | 0.284 | 2.255 |
| Russian | Han, Mbuti, Karitiana, Ulchi and Mixe | 0.042 | 0.517 | 0.441 | 5.389 |
| Russian West | Han, Mbuti, Karitiana, Ulchi and Mixe | 0.035 | 0.526 | 0.440 | 0.819 |
| Croats | Han, Mbuti, Karitiana, Ulchi and Mixe | 0.042 | 0.303 | 0.655 | 4.373 |
| Germans | Han, Mbuti, Karitiana, Ulchi and Mixe | 0.027 | 0.373 | 0.6 | 5.999 |
| Poles | Han, Mbuti, Karitiana, Ulchi and Mixe | 0.023 | 0.561 | 0.416 | 2.634 |
